# Photobiomodulation Therapy for a Novel Olfactory Dysfunction Ischemic Stroke Model

**DOI:** 10.1101/2023.02.07.527573

**Authors:** Reham. A Shalaby, Muhammad Mohsin Qureshi, Mohd. Afzal Khan, S. M. Abdus Salam, Hyuk Sang Kwon, Kyung Hwa Lee, Euiheon Chung, Young Ro Kim

## Abstract

A.

**Background:** Ischemic stroke typically accompanies numerous disorders ranging from somatosensory dysfunction to cognitive impairments, inflicting its patients with various neurologic symptoms. Among pathologic outcomes, post-stroke olfactory dysfunction is frequently observed. Despite the well-known prevalence, therapy options for such compromised olfaction are limited, likely due to the complexity of the olfactory bulb architecture, which encompasses both the peripheral and central nervous systems. As photobiomodulation (PBM) emerged for treating stroke-associated symptoms, the effectiveness of PBM on the stroke-induced impairment of the olfactory function was explored.

**Purpose:** To address the efficacy of PBM therapy on the olfactory bulb damage caused by ischemic stroke using both behavioral and histologic and inflammatory markers in the newly developed stroke mouse models.

**Methods:** Novel mouse models with olfactory dysfunction were prepared using photothrombosis (PT) in the olfactory bulb on day 0. Moreover, post-PT PBM was performed daily from day 2 to day 7 by irradiating the olfactory bulb using an 808 nm laser with the fluence of 40 J/cm^2^ (325 mW/cm^2^ for 2 minutes per day). The buried food test (BFT) was used for scoring behavioral acuity in the food-deprived mice to assess the olfactory function before PT, after PT, and after PBM. Histopathological examinations and cytokine assays were performed on the mouse brains harvested on day 8.

**Results:** The results from BFT were specific to the individual, with positive correlations between the baseline latency time measured before PT and alterations at the ensuing stages for both the PT and PT+PBM groups. Also in both groups, the correlation analysis showed a significant positive relationship between the early and late latency time changes independent of PBM, implicating a common recovery mechanism. In particular, the PBM treatment largely accelerated the recovery of impaired olfaction after PT with the suppression of inflammatory cytokines while enhancing both the glial and vascular factors (e.g., GFAP, IBA-1, and CD31).

**Conclusions:** The PBM therapy during the acute phase of ischemia improves the compromised olfactory function by modulating the microenvuronment and tissue inflammation.

## B. Introduction

Photobiomodulation (PBM) is a drug-free light therapy that is well-documented for its beneficial effects via endogenous self-repair mechanisms [1]. Numerous studies have demonstrated the therapeutic efficacy of PBM, which spans across a wide range of measures in both structural and functional recoveries. The PBM therapy-associated biological alterations include the reduction of inflammation and oxidative stress, as well as the improvement of mitochondrial function [2-4]. Moreover, previous clinical reports also indicated treatment benefits by multiple exposures, such as the stimulation of wound healing [5], pain alleviation in various orthopedic and musculoskeletal conditions [6], and restoration of brain function [7].

From the neurological aspect, the efficacy of PBM has been shown in treating the traumatic brain injury with outcomes including increases in the cerebral volume and perfusion, and improvements in the functional connectivity and cognitive faculties [8, 9]. PBM is also known to be effective in the management of patients with Alzheimer’s dementia by boosting executive function, memory, visual attention, and connectivity measures [10, 11]. Additionally, preclinical studies revealed that the PBM therapy could be also effective in Parkinson’s disease [12] and ischemic stroke [13, 14]. In particular, previous stroke model studies revealed that the PBM therapy reduces behavioral deficits while promoting neurogenesis [13]. Although emerging as one of promising candidates for treating stroke-associated functional impairments, mechanisms of action underlying such recovery have been elusive.

In this regard, using a novel animal stroke model with function-specific damage, our current study aims to address the utility of PBM by demonstrating the direct behavioral/functional recovery in conjunction with histopathological assays. We targeted the olfactory bulb (OB) as our stroke site of interest considering the profound prevalence of post-stroke olfactory dysfunction, which is present in 43.6 % of the stroke patients, a year after the insult [15]. Additionally, such olfactory dysfunction is also commonly observed in patients with Alzheimer’s disease [16], Parkinson’s disease [17], and frontotemporal dementia [18]. Reports also suggested that the impaired olfaction is linked with cognitive decline and might be used as an indicator of neurodegeneration among dementia-free older adults [19]. More recently, pre-symptomatic and aftereffect olfactory dysfunctions in Covid-19 patients have been recognized as common traits of the disease, further augmenting the need for developing effective therapeutic strategies to treat the olfaction related impairments [20].

To induce a olfactory function loss via highly controlled formation of the ischemic stroke, we adopted photothrombosis (PT) precisely at the OB to which the localized PBM therapy was directly applied daily., Following the PT induction, behavioral measurements, immunohistochemistry, and cytokine assays were performed to explore the factors involved with functional changes. The importance of our current work lies in the experimental demonstration to answer whether the PBM therapy influences the recovery of impaired olfactory function while affecting the associated biological/biochemical factors. Positive outcome of the proposed non-invasive PBM approach would strongly benefit patients suffering from olfactory symptoms derived from stroke and other neurodegenerative diseases.

## C. Methods

All experimental protocols followed the guidelines of the Institutional Animal Care and Use Committee (IACUC) at Gwangju Institute of Science and Technology (GIST), Korea. The experimental protocols were approved by the GIST IACUC under protocol # GIST-2020-077. We used 66 C57BL/6 male mice aged between 10 and 12 weeks. All mice were 25-30 grams in body weight.

### a. Buried food test (BFT) for olfactory function measurement

The buried food test (BFT) was performed according to the previously published protocol [21]. Adaptation to food was achieved by introducing the food pellets with fresh odor daily to the mice for one week. On the test day, animals were fasted (except for water) for 18 hours and then introduced to a clean mouse cage where an odorous food pellet (RodFeed, DBL, Chungcheongbuk-do, South Korea) was randomly embedded under the 3 cm thickness of clean bedding. Then, latency in the food search time was measured, which was repeated three times for each BFT session. The first BFT after PT was performed three days after the PT since certain mice were inactive after their PT surgeries. So the time was taken to allow the mice to recover, as shown in Figs 1A and 1B. We measured the body weight before each BFT to assess the weight fluctuation after 18 hours of fasting. The BFT was conducted on day - 1 before the PT induction to obtain the baseline which was followed by the BFTs on day 3 and 7 post-PT.

**Figure 1.**
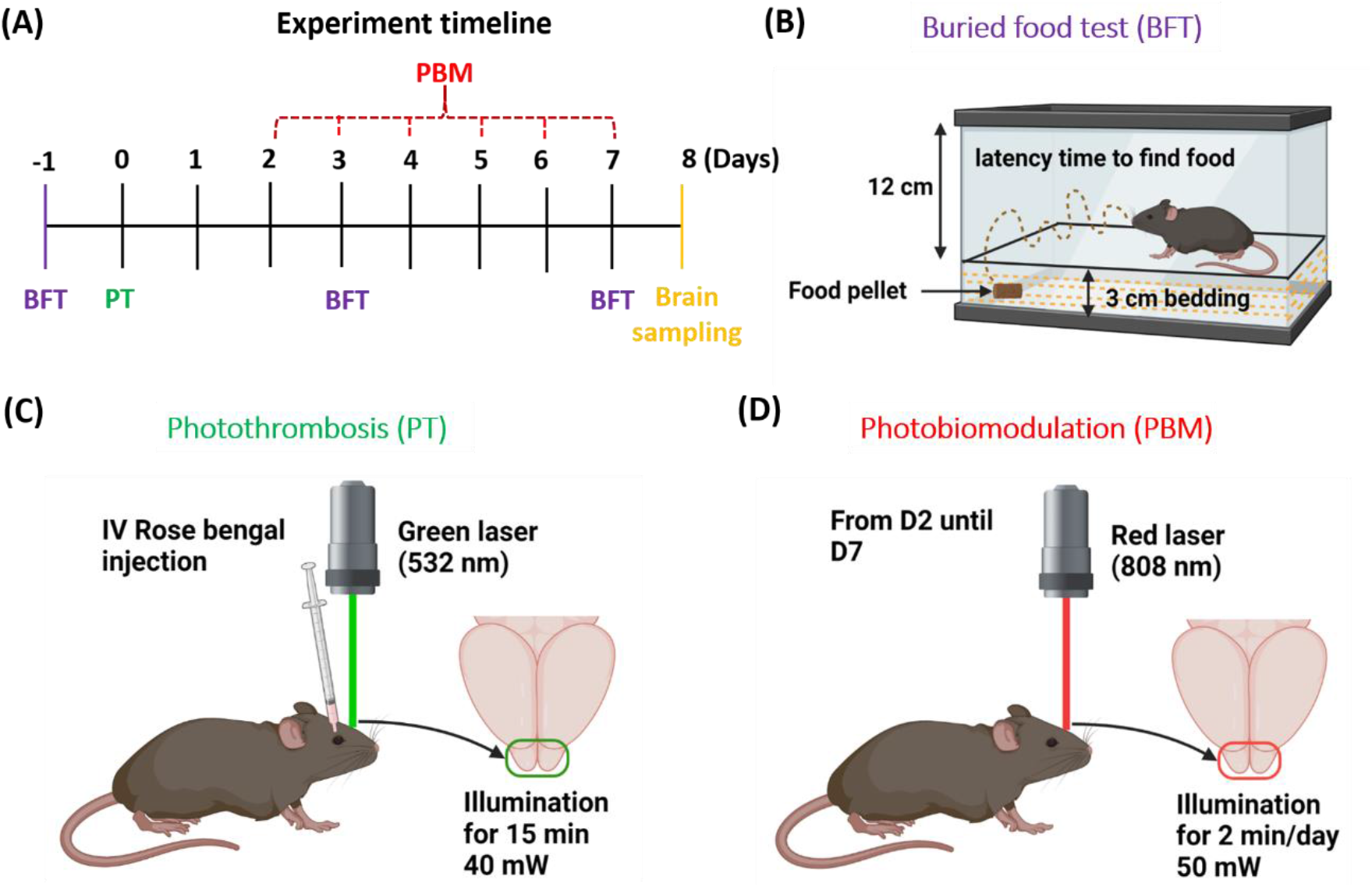
Schematic diagram of experimental design. (A) Experimental timeline of olfactory function test (i.e., buried food test: BFT), photothrombosis (PT), photobiomodulation (PBM), and brain sampling. (B) Illustration of BFT. (C) Photothrombosis parameters using a 532 nm green laser. (D) Photobiomodulation parameters using an 808 nm red laser.

### b. Olfactory bulb (OB) photothrombosis

Photothrombosis (PT) was performed on day 0 for both PT and PT+PBM groups. An incision was made in the scalp over the OB area between two eyelids; the periosteum was gently removed using a scalpel. Then a 4 mm coverslip was fixed over the OB area and sealed with glue. A retro-orbital injection of 20 mg/kg of rose Bengal was given to the PT group, after which a green 532 nm laser was illuminated over the coverslip for 15 minutes in a dark room, as shown in Fig 1C. Then, each mouse was transferred to a new cage and placed for recovery. The identical protocol was used for the control group, except for the rose Bengal injection. The body temperature was maintained at 37 ± 0.5°C throughout the procedures.

### c. Photobiomodulation after photothrombosis in olfactory bulb

Photobiomodulation was performed daily using an 808 nm laser from day 2 to 7 post-PT for 2 minutes per day. The fluence was 40 J/cm^2^ per session, where the laser beam diameter was 4.17 ± 0.07 mm with the irradiance of 325 mW/cm^2^, as shown in Fig 1D. The PBM was performed in a dark room while maintaining the body temperature at 37 ± 0.5°C. A holed black mask was used over the OB area to protect the surrounding tissue from unnecessary light exposures.

### d. Photobiomodulation in sham control mice

The PBM control group was prepared with the same surgery and by placing the coverslip over the OB but without inducing the photothrombotic ischemia (i.e., excluding the illumination of a 532 nm laser and injection of rose bengal). Using an 808 nm laser, PBM was similarly applied in this control group from day 2 to 7 (i.e., Sham Control +PBM), while performing the BFT to evaluate functional changes.

### e. Brain sampling

On day 8, mouse brains were harvested after the heart perfusion with 4% paraformaldehyde solution (PFA) and fixed in 10% neutral-buffered formalin for three days. Then, the brains were dissected, embedded in paraffin, and stained with hematoxylin and eosin (H&E) for histologic examination. For the cytokine assay, brains were extracted on day 8 without the perfusion and immediately kept in a -80°C freezer.

### f. Cytokine assay

The post-PT cytokine levels were assessed on day 8 using the OB tissue lysates obtained from the control, PT, and PT+PBM groups. The cytokines expressions were analyzed by Proteome Profiler Mouse Cytokine Array Panel A Kit (ARY006, R&D Systems) on 9 samples (3 from each group). Expression levels were analyzed densitometrically using Image J, after being corrected by the values determined in positive controls and expressed as relative values compared to the control group.

### g. Immunohistochemistry

Paraffin blocks of mouse brains were cut in 3 μm thickness and subjected to immunohistochemistry (IHC). Tissue sections were stained with antibodies against Glial Fibrillary Acidic Protein (GFAP) (dilution 1:400, code no. M0761, Dako, Glostrup, Denmark), IBA1 (dilution 1:400, code no. 234-003, Synaptic Systems, Goettingen, Germany) and CD31 (dilution 1:200, code no. M0823, Dako) using the Bond-max system (Leica Microsystems, Bannockburn, IL, USA). Negative controls were processed similarly in the absence of primary antibodies. The resulting slides were evaluated by a clinical pathologist (KHL), which were digitalized using a slide scanner (Aperio ScanScope CS System, Leica Microsystems) and photographed by the Aperio Image Scope 12.3 (Leica). For the analysis of IHC, three ROI’s in the stained image were selected for each OB at ∼200 μm from the lesion, corresponding to the area surrounding the lesion (i.e., peri-infarct area). To analyze images of GFAP and IBA-1, Image J (Fiji) software was used for the quantitative evaluation of immunoreactivity. The images were inverted using Image J, and the numbers of immune-positive cells were counted with a multi-point tool. For the CD31 quantification, automated analyses were performed to count the number and mean area of the vessels using a homemade algorithm and the open CV library.

### h. Statistical analysis

Statistical analysis for determining group differences was performed using one-way analysis of variance (ANOVA) tool in the GraphPad Prism software. Student’s t-tests were used when two groups were compared for the difference. Statistical significance was accepted at the 95% confidence level (P < 0.05). Data were expressed as means ± SD.

## D. Results

### a. Development of olfactory bulb photothrombosis model

We developed a novel rodent model of OB ischemia based on PT using a 532 nm green laser, as shown in Fig 2. Since the green laser beam profile is directly linked with the formation of ischemic damage, the spatial laser beam profile was characterized and tested for repeatability. The diameter of the 532 nm laser beam used in our study was 3.9 ± 0.1 mm. In order to quantify the olfactory function and to explore the ideal time of intervention with PBM and also the proper time for brain sampling, we utilized the BFT and measured the latency times on the day before PT (D-1), day 3 (D3), D7, and D14 after PT. As shown in Fig 2C, the latency time (i.e., the time taken before the discovery of hidden food) significantly increased on D3 compared to the control value before PT (i.e., D-1) for the PT group, which was slightly but significantly reduced on D7 when compared to D3. Between the D7 and D14, no further change in the latency time was observed. The H & E staining provided the detailed description of the tissue alterations after PT, revealing an abnormal tissue morphology with degenerative features, in contrast to those found in the normal controls as shown in Fig 2D.

**Figure 2.**
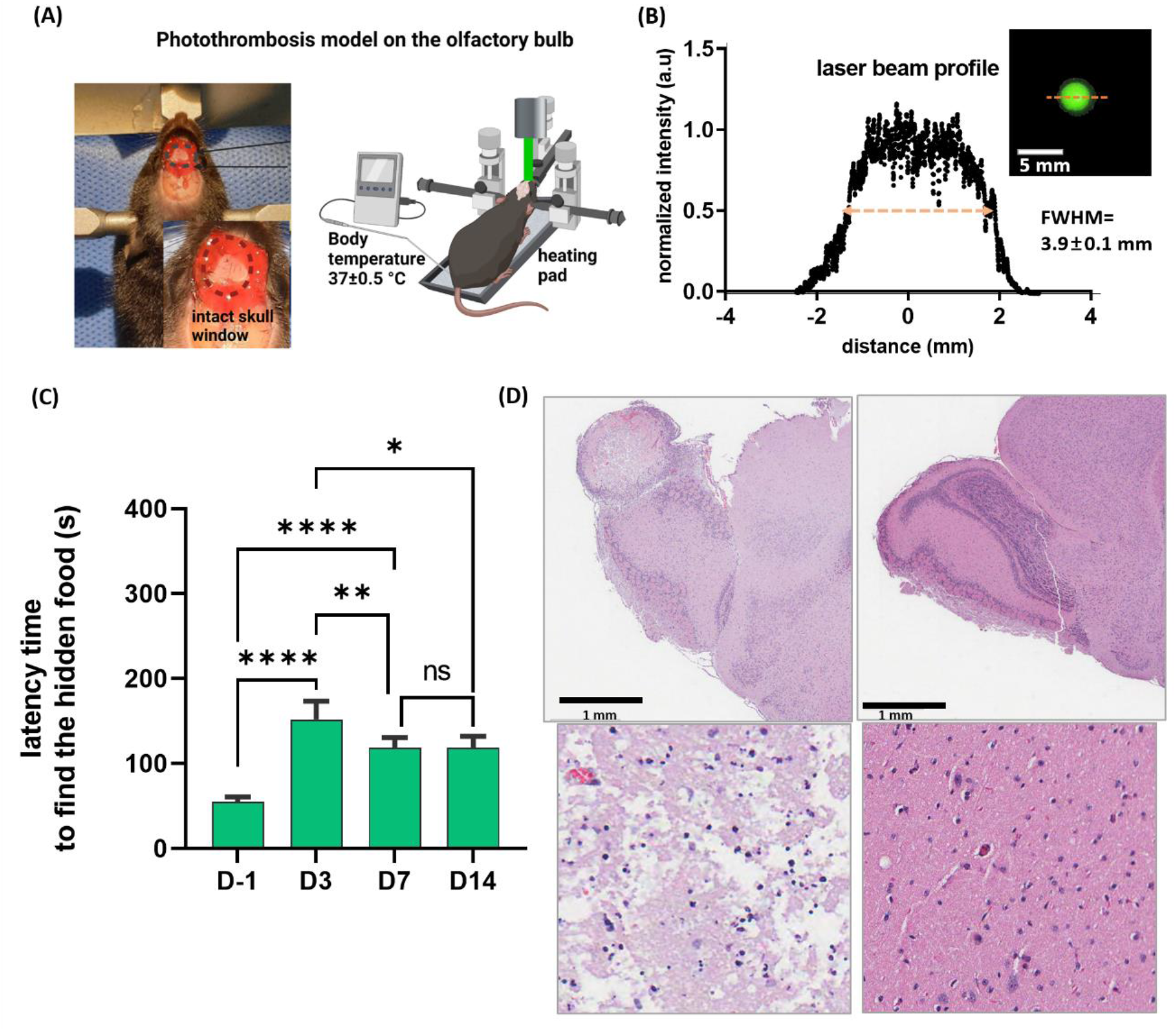
Olfactory bulb photothrombosis model. (A) Phtothrombosis setup and parameters. (B) 532 nm laser beam profile during OB illumination. (C) Olfactory function test results after photothrombosis, n=5. (D)H&E staining images of the post-PT OB with high magnification, taken at 20 X showing degenerative features, compared with the sham control OB with high magnification. Data are mean ± SD. (n=5; *P<0.05, **P<0.01, ****P <0.0001; ns-not significant.

### b. PBM significantly improves behavioral deficits after PT

As shown in Fig 3, in normal control (NC) mice, the latency time did not vary throughout the test days, confirming the reference stability of BFT over time. For the sham control (SC) group, the olfactory function was unaffected after the 532 nm laser exposure as also, no PBM-related alterations were found in the normal, healthy olfactory function of the SC+PBM mice. The findings reveal that the laser exposures caused little influence on the normal olfaction, in which PBM using an 808 nm laser was applied between D2 and D7 after surgery for 2 minutes per day.

**Figure 3.**
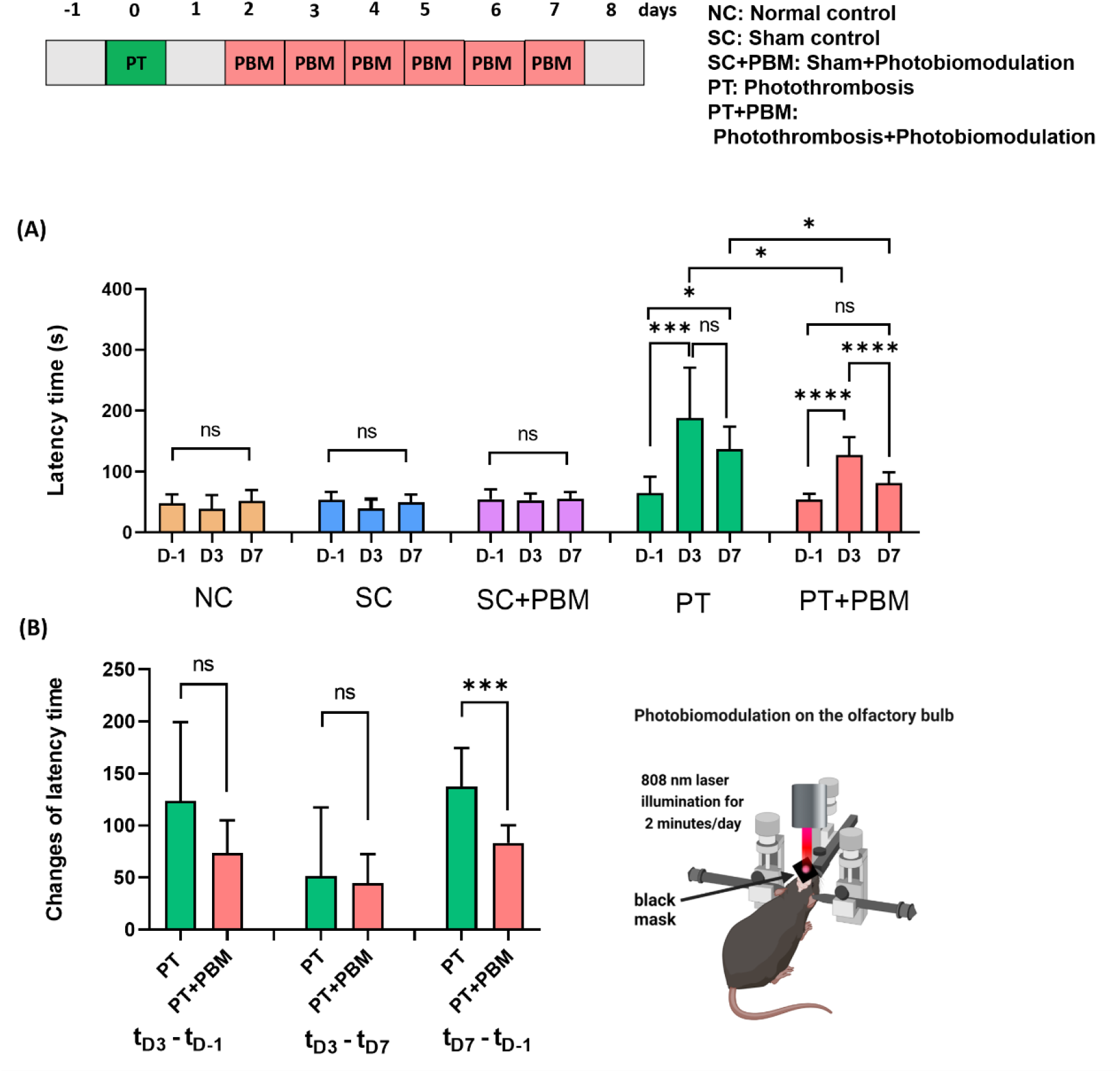
Effects of PBM on behavioral deficits after PT. (A) Buried food test results for all experimental groups over time, NC: n=8, SC: n=8, SC+PBM: n=6, PT: n=11, PT+PBM: n=10,. (B) Olfactory function test results from the changes of the latency times for the PT and PT+PBM groups. Data are mean ± SD. (*P<0.05, ***P<0.001, ****P <0.0001; ns-not significant).

The BFT’s were performed on D-1, D3, and D7 to investigate the impact of PBM following PT. The latency time measured on D3 for the PT mice significantly increased compared to that of the baseline (D-1). In comparison with the PT group, the PT+PBM group exhibited a significant improvement in the olfactory function on D3. For the PT mice, the significantly deteriorated olfactory function on D3 only improved with a slight and non-significant latency time change between D3 to D7 as in Fig 3A. Of a note, the latency time on D7 was significantly longer than that of the baseline (D-1) in the PT group whereas these values were similar for PT+PBM. Upon comparison with the PT group, significantly greater improvements of the impaired olfactory function were observed on both D3 and D7 in the PT+PBM group. In particular, the significantly improved olfactory function on D3 revealed a clear, positive effect even by a single, one-time PBM treatment (see the comparison between PT and PT+PBM groups in Fig 3A).

As shown in Fig 3B, a quantitative analysis revealed that the latency time changes from the baselines were significantly different between the PT and PT+PBM groups only on D7 (after six sessions of PBM). Such group differences of the latency time changes were not pronounced in other periods (i.e., t_D3_-t_D-1_ and t_D3_-t_D7_), implicating the efficacy of the long term PBM therapy. In addition, it was notable that for both PT and PT+PBM groups, the outcomes of BFT were specific and able to delineate the individual differences, as the significant positive correlation was found between the baseline and changes in the latency time (see Fig 3B and supplementary Fig S3).

### c. Correlation analysis of the latency time changes

Correlation analyses among the latency time changes revealed that there were significant, a positively correlative relationship in the changes of the latency time between the early (D-1 and D3) and late (D3 and D7) periods as shown in Fig 4 (A) and (B). No other significant relationships were found upon analyzing the correlation among the latency times and baselines.

**Figure 4.**
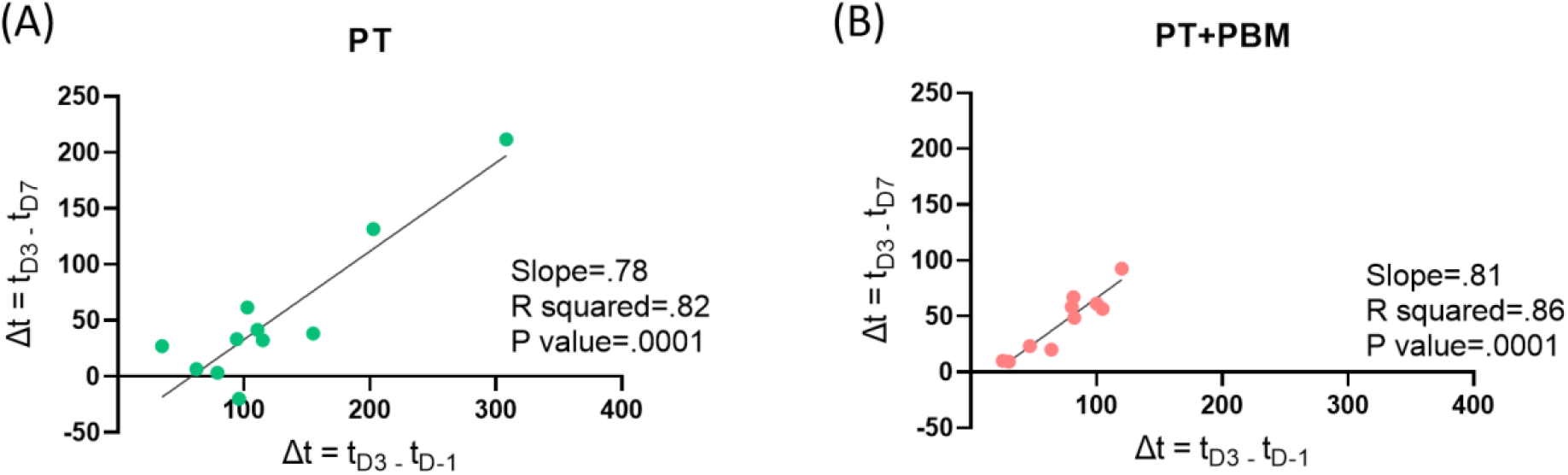
Correlation of changes in the latency time between the early (D-1 and D3) and late (D3 and D7) periods for (A) PT and (B) PT+PBM groups.

### d. PBM has anti-inflammatory effects after PT

As shown in Fig 5, higher expression levels of inflammatory cytokines (e.g., C5/C5a, IL-1α, IL-1β, IL-16, etc.) were detected in the PT group when compared to the control group. As demonstrated by the PT+PBM group, such inflammatory cytokine levels tended to decrease by PBM. For certain cytokines (e.g., IL-16, IL-17, IFN-γ, TNF-α), the expression levels were restored by PBM near to the control levels. The anti-inflammatory mediators were not altered by PT but responded to PBM, in which cytokines including GM-CSF, IL-1ra, IL-4 and IL-10 were up-regulated to the levels higher than those from the control group (Figs 5C). In addition, we also found that sICAM-1, the soluble form of cellular adhesion molecule, a useful marker of endothelial activation and inflammation was highly expressed after PBM (Figs 5A). One-way ANOVA was used to evaluate the statistical significance between the three groups. Thus, PBM did not only inhibit the post-PT inflammatory activities but also promoted anti-inflammatory effects.

**Figure 5.**
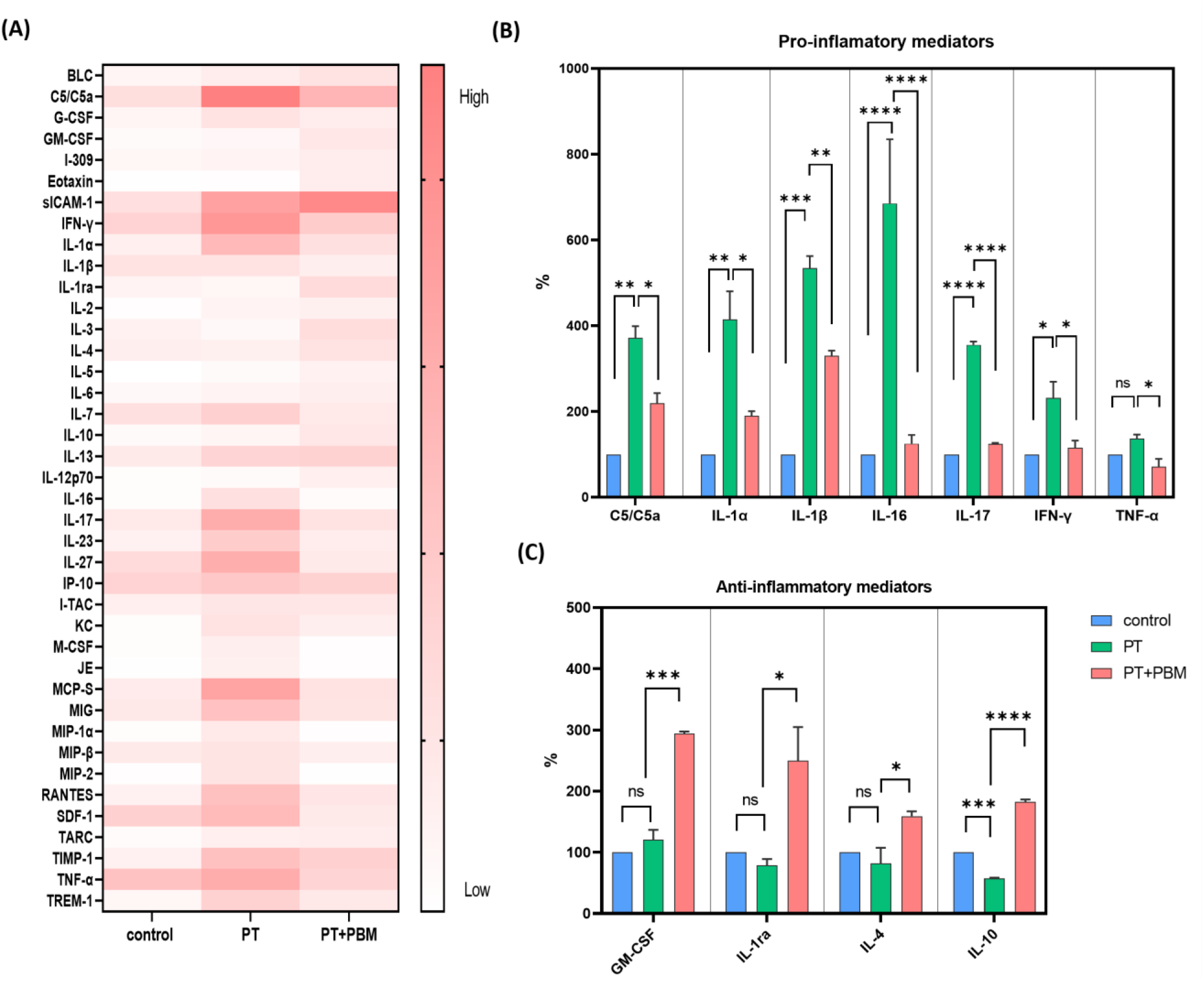
The effect of PBM on cytokines after PT. (A) Heat map visualization of cytokines and chemokines for control, PT and PT+PM mice. (B) Effects of PT and PBM on the pro-inflammatory and (C) anti-inflammatory mediators. n=3/group.) Mean ± SD. (*P<0.05, **P<0.01, ***P<0.001, ****P <0.0001; ns-not significant.

### e. PBM enhances microenvironment recovery after PT

As shown in Fig 6A-6D, glial fibrillary acidic protein (GFAP: present mostly in reactive astrocytes) expression was significantly higher in the PT group than either the control or the PT+PBM group measurements, implicating the suppression of the post-PT astrocyte activation by PBM. In Fig 6E–6H, the expression of allograft inflammatory factor 1 (IBA-1), responsible for injury repair was significantly higher in both PT and PT+PBM groups than control. Such trend suggested the highly enhanced post-PT activation of microglia, which was unhampered by the PBM treatment. As shown in Fig 7A-7C, our results additionally demonstrated that the stains of CD31 expression in the PT+PBM group was more densely populated than that found in the PT group. And both the microvascular density and demarcated area were significantly higher in the PT+PBM group than PT, exhibiting post-PT restoration of the microvascular matrix by PBM.

**Figure 6.**
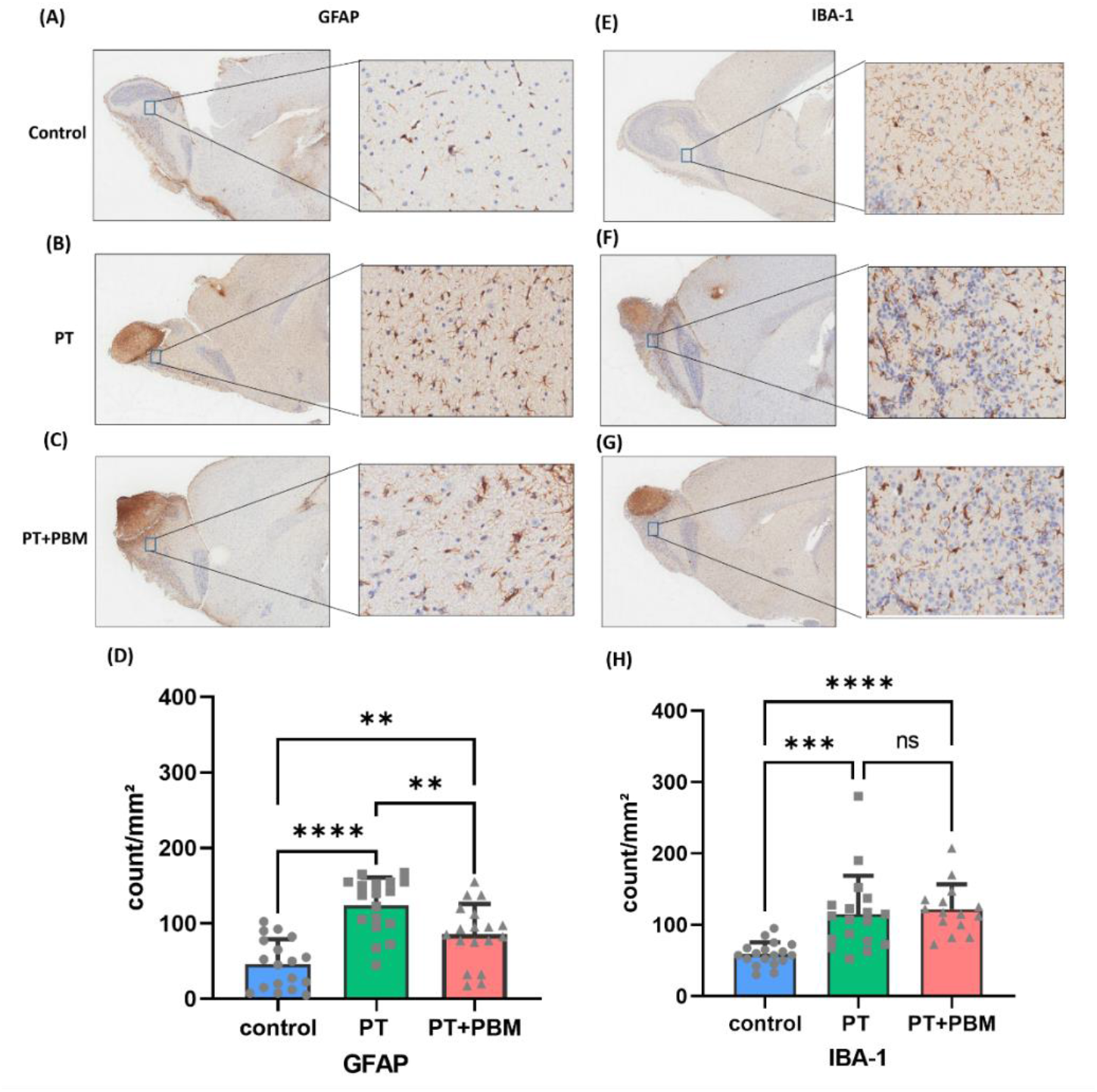
PBM manipulates the activation of non-neuronal cells after PT. (A-C) Expression of GFAP in control, PT, PT+PBM with high magnification images. (D) Quantitative analysis of GFAP. (E-G) Expression of IBA-1 in control, PT, PT+PBM. (H) Quantitative analysis of IBA-1. n=6/group. Data are expressed as mean ± SD. (n=6; *P<0.05, **P<0.01, ***P<0.001, ****P <0.0001; ns-not significant.

**Figure 7.**
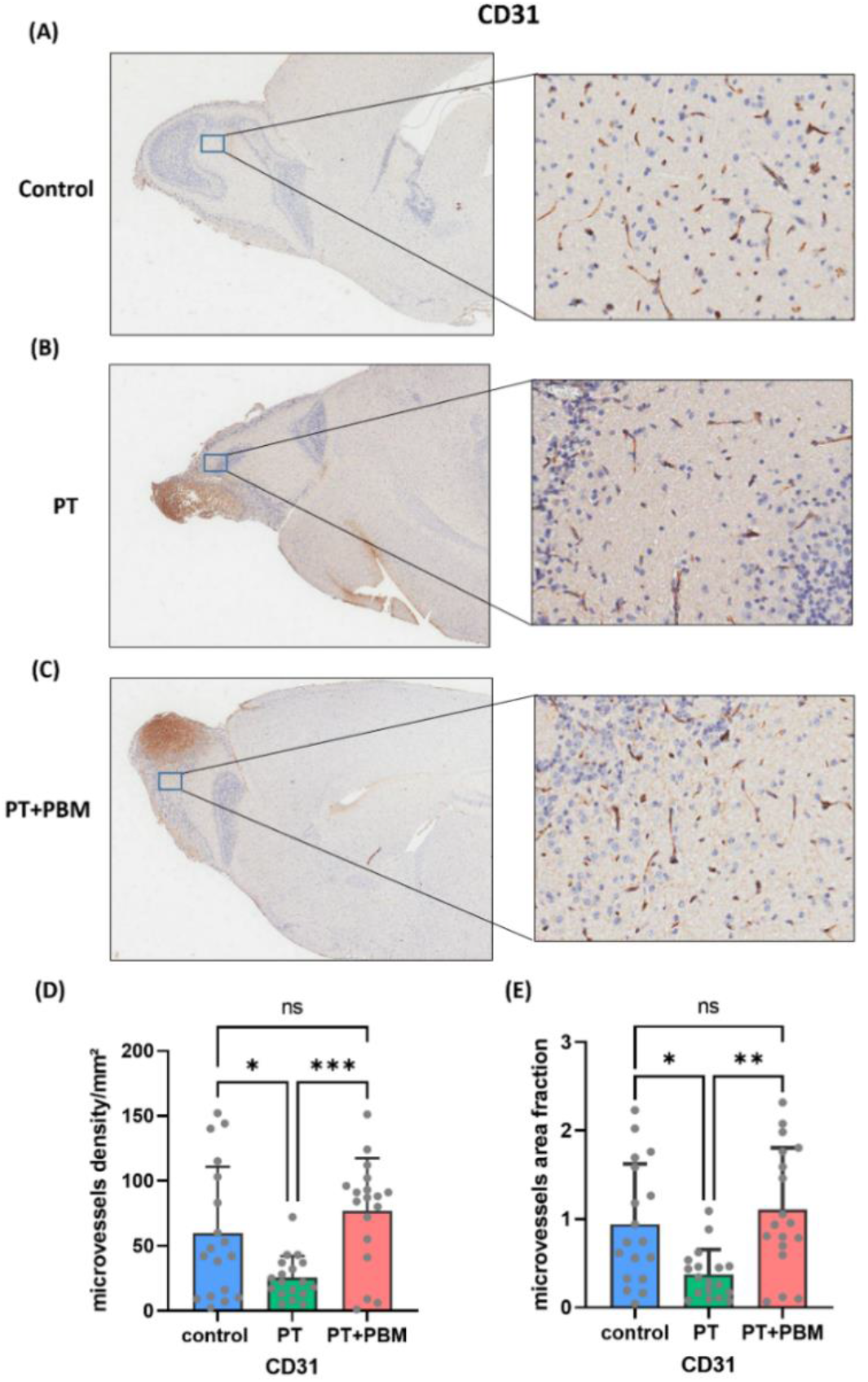
PBM enhances the expression of CD31 after PT. (A-C) expression of CD31 in control, PT, PT+PBM with high magnification image. (D) Quantification of CD31count for control, PT, and PT+PBM. (E)Quantification of the CD31 area. Data are mean ± SD. (n=6; *P<0.05, **P<0.01, ***P<0.001; ns-not significant. n=6/group.

## E. Discussion

The current study was designed to explore the therapeutic efficacy of photomodulation for treating olfactory dysfunctions. One of the primary aims in our study was to investigate the effects of PBM on neurofunctional impairments using an animal model that mimics human brain disease conditions. In this regard, we developed a new animal model by precluding the use of any unnatural intervention, such as genetic engineering or chemical modifications, which are common in Parkinson’s and Alzheimer’s disease [22, 23]. For this purpose, we chose to utilize focal ischemia, which selectively disrupts the neural function via occlusion of the local blood flow.

In particular, to increase the specificity of the functional deficit, we induced photothrombosis (PT), which was regionally confined in the olfactory bulb of mouse. Highly precise for damaging the brain area of interest, it is also notable that the PT-induced stroke produces a relatively small ischemic penumbra [24]. Since penumbral peri-infarct areas are known to be involved in the recovery process, the lack of such might have limited the current study in the inexact pathological reproduction of human stroke conditions. However, compared with other traditional stroke methods, such as filament occlusion, the PT-induced lesions were highly reproducible for maintaining the consistent infarct size and location, only impairing the olfactory function as proposed in our study. Such specificity of PT was highly important to confine the study focus concentrated on the examination of factors that are strictly associated with the olfaction behavior.

As for assessing the functional deficit, buried food test (BFT) has been used frequently to quantify behavioral acuity of the olfactory function [25]. While there is a controversy due to the possible reliance on locational memory, in the current study, such shortcoming was mitigated by randomly placing the food pellet and also by measuring the latency time three times at each trial. A few previous studies have indicated that even a single dose of photonic therapy could confer beneficial effects on brain injuries, independent of the therapy time point after the disease onset. Similarly, in the current PT model, functional improvements were observed by a single PBM treatment on post-PT D3 (see Fig 3A), in agreement with reports from previous studies [26-29]. Moreover, by tracking down the changes in individual performance over the course of post-PT measurements (e.g., excluding the baseline contribution), we were able to delineate subtle functional alterations. Upon comparing the PT group with PT+PBM based on such approaches, effects of the PBM treatments were demonstrated with a particular functional improvement on D7, which was not observed on D3 as shown in Fig 3B. The significantly improved performance on D7 shows that the repeated PBM treatments are likely more effective than a single treatment [13, 30]. Nonetheless, future studies are needed for determining the exact therapy regimen to achieve optimal effectiveness of the PBM treatment.

Interestingly, the changes in the latency time, indicated that the rate of function shifts at early stages predicted the recovery rate at the late stages. As shown in Fig 4, we found out that the significant correlations, independent of the PBM treatments were present between the functional changes quantified over the D-1∼D3 and D3∼D7 periods. More Remarkable, the correlation slopes were similar between the PT and PT+PBM groups. Such similarity indicates a possibility of the recovery mechanism shared by both the PT and PT+PBM groups, in which the PBM treatments appeared to accelerate the recovery process.

During the pathological progression of ischemic stroke, management of the acute inflammatory response plays an essential role in the prognosis of patient outcomes. Although neuroinflammation, in general provides a defense mechanism that removes cell debris and promotes tissue repair at the initial stage of stroke progression, the prolonged presence of inflammatory factors could exert an adverse effect on the functional recovery [31]. Therefore, we posited that the attenuation of the pro-inflammatory cytokines is likely associated with protective mechanisms in the post-ischemic brain tissue [32]. As demonstrated in Fig 5, PBM not only downregulated the inflammatory cytokines after PT but also upregulated the levels of anti-inflammatory cytokines. Such dynamic levels of cytokines could play a vital role in the pathogenesis of stroke, which reciprocally poses as potential targets of therapeutic intervention [33]. Although the mechanisms are not yet well defined, we showed that the elevated expression of pro-inflammatory cytokines such as IL-1α, IL-1β, IL-16 after PT was significantly reduced by PBM. Also important, the levels of anti-inflammatory cytokines including IL-1ra, IL-4 and IL-10 were effectively elevated in the PT animals treated with PBM compared with the PT group [13]. In this regard, our study showed that PBM reduces further deterioration of the olfactory function by inhibiting the inflammatory process as well as by enhancing the healing process by increasing the blood flow leading to tissue oxygenation and enhancing neurogenesis after stroke.

In addition, the upregulated CD31 demonstrated that PBM enhances the restorative blood vessel density in the olfactory bulb after PT, likely promoting the circulation-related recovery process [34]. On the other hand, the reactive astrocytes (delineated by GFAP: see Fig 6), which were significantly reduced in the PT+PBM group, can be functionally divided into both neuroinflammatory and neuroprotective types (A1 and A2) [35]. In contrary to the PBM-induced restoration of olfactory function, the neuroinflammatory type (A1) is known to possess a significant neurotoxicity after stroke and is associated with the worsening neurological prognosis [36]. We hypothesize that PBM selectively inhibits the activation of toxic astrocytes, for which future studies are warranted.

We also demonstrated that the elevated inflammatory mediators such as C5a, IL-1α, IL-1β, IL-16, IL17and TNF-α which are known to promote neuronal apoptosis, aggravate brain injury, inhibit neurogenesis, promote adhesion molecules, and reduce the blood-brain barrier integrity [37-39] were significantly downregulated by PBM. Moreover, PBM highly enhanced the anti-inflammatory mediators like GM-CSF, IL-1ra, IL-4, and IL-10, previously reported to have neuroprotective effects by reducing inflammatory markers associated with risks of poor outcomes, thus promoting the stroke recovery [13, 40]. On the contrary, also highly affected by PBM was the elevated level of sICAM-1 which is known to correlate with cerebral microbleeds and hemorrhagic transformation in stroke patients [41] participating in the deterioration of the post-stroke vascular function. For resolving such discrepancy, focused future studies are required to address the role of sICAM-1 in the post-stroke functional recovery and to reveal the mechanism of PBM in its expression.

To date, most animal stroke studies have focused on the cortical and hippocampal regions of the brain, which are associated with multiple dysfunctions encompassing learning, memory, motor, etc. Although the importance of the olfactory function and its implication in many neurodegenerative diseases has been reported frequently, there are only few lesion studies conducted directly on the olfactory bulb. In our rodent study, a novel window model exposing the olfactory bulb allowed an easy access for the optical *in vivo* modulation, while the effect of damage could be quantified by the olfactory function assessment test (e.g., BFT). From the translational perspective, future studies can be designed to optimize a transcranial PBM strategy for the post-stroke patients with a loss of olfactory function by employing transcranial near-infrared light helmets. Meanwhile, PBM can be also administered through the nasal cavity and stimulate the recovery process [42], thus making it a potential therapeutic option for COVID-19 patients who are prone to have olfactory dysfunction with the local tissue damage [43-45].

Although a few inflammatory factors were investigated in the current study, specific mechanisms underlying the functional recovery after PBM are yet to be determined. Nonetheless, based on the current findings, we concluded that the selective expressions of genes that have key roles in the stroke recovery process are promoted by PBM rather than the restoration of the infarcted tissue. The importance of our study lies in the demonstration of accelerated functional recovery by the PBM therapy. Moreover, such functional restoration can be emphasized for conducting feasibility investigations of PBM as a clinical therapy option to treat patients with various olfactory functional deficits.

## F. Conclusion

Using photothrombosis (PT) in the olfactory bulb of mouse brain, we developed a new ischemic stroke model that is optically accessible and evaluated the efficacy of photobiomodulation (PBM) for treating the impaired olfaction behavior. Specifically, we showed that the daily PBM therapy could significantly improve the compromised olfactory function while exerting little effects on the normal healthy brains. These outcomes were additionally supported by the histopathological staining and cytokines assay, revealing the role of PBM for suppressing post-stroke inflammatory reactions. As a result, we PBM could be a promising therapy option at the acute stage of brain ischemia to accelerate the functional recovery process and possibly, for treating olfactory dysfunctions resulting from various etiological causes including COVID-19.

## Supporting information

supplemetary materials for the experimental results.

## G. Acknowledgment

The authors thank Eun Jung Ahn for assisting in the Cytokine assay, Minsung Kim for helpful discussion, Akm Ashiquzzaman for assisting in CD31 analysis and Christine H. Hwang for reviewing the manuscript.

