## Supplementary material for "Photobiomodulation Therapy for a Novel Olfactory Dysfunction Ischemic Stroke Model": supplemetary materials for the experimental results.

### Supplementary materials

#### a. PBM has a safe thermal effect on animals

To assess the thermal effect of an 808 nm laser on the animal brain, we measured the temperature changes before, during, and after illuminating the laser. This helped us to find that the temperature takes 30 seconds to increase by 5~6℃ and become stable. The temperature also took 30 seconds to return to its normal level. The temperature did not rise over 40 ℃, indicating that the current laser illumination is safe and would not cause brain damage, as shown in Figure S1 (B). Moreover, the animals showed no abnormal behavior after exposure to PBM. The 808 nm laser beam profile was measured and analyzed using ImageJ software, as shown in figure S1 (C).

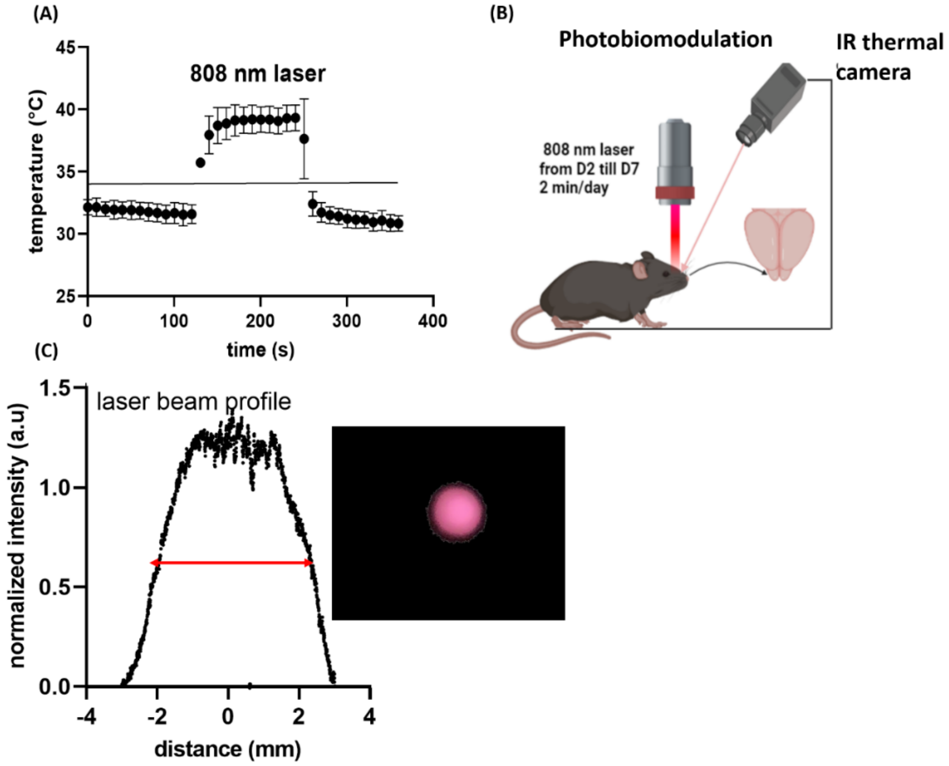


Fig S1. **Temperature measures during PBM.** (A) Temperature fluctuations during 808 laser illumination. (B) Schematic diagram of PBM and its temperature fluctuations. The temperature was measured 2 minutes before and 2 minutes after illuminating the laser. (C) The beam profile from five trials of illumination, FWHM (full-width-at-half-maximum) is about $4.17\pm0.07$ mm.

**b. Weight changes after fasting in PT and PT+PBM groups**

We measured the weight of each experimental animal after fasting them to determine if fasting would significantly affect their weight. We found that the mice lost about 5% of their weight during fasting for 18 hours. However, after fasting, the mice started to gain weight again. Thus, we concluded that PT and PBM do not affect the mice's weight (Figure S2).

**c. The relation between baseline and impaired olfactory function**

In order to determine the relationship between the baseline olfactory function at D-1 and the impairment at D7, the paired comparison was plotted. It was found that the majority of the mice exhibit the same trend, with mice with short baseline latency times having shorter latency times after PT and vice versa (Fig S3).

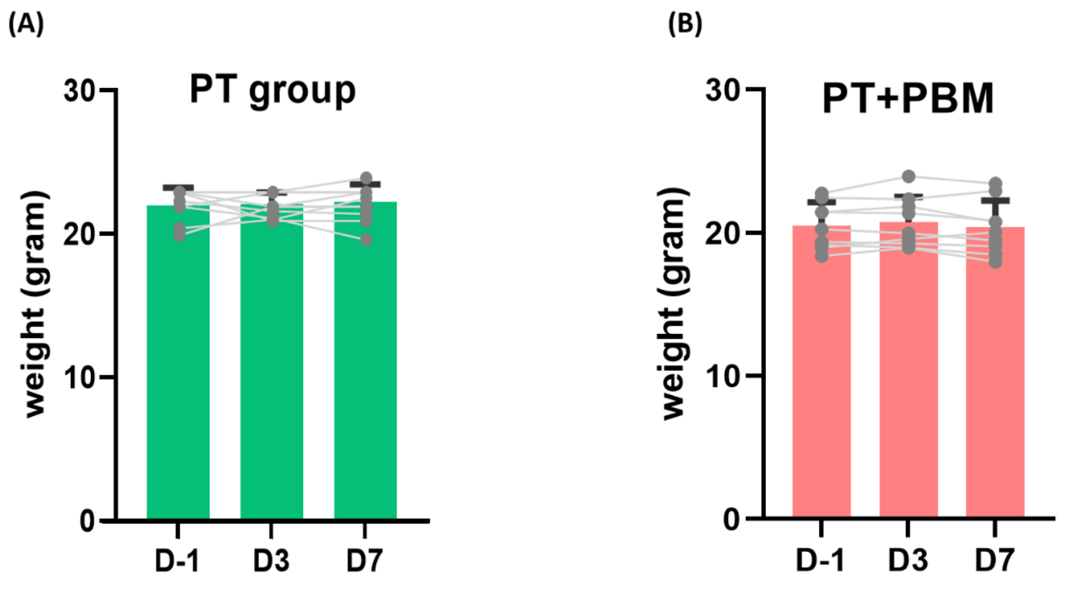


Fig S2. **Weight fluctuations over time**. (A) Weight fluctuations in the PT group. (B) Weight fluctuations in the PT+PBM group.

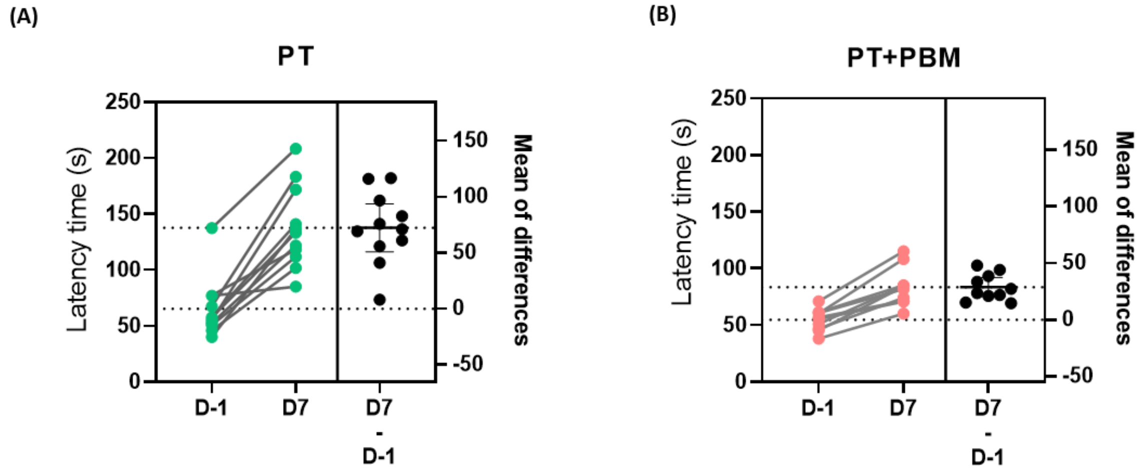


Fig S3. **Relationship between baseline and impaired olfactory function** (A), (B) The paired baseline latency time with the impaired one for PT and PT+PBM groups (The mean latency times for before (D-1) and after (D7) are indicated as dotted horizontal lines: the lower line for D-1 and the upper line for D7)

#### d. PBM accelerates the natural recovery after PT

According to the severity of the olfactory dysfunction, the olfactory function impairment was categorized as heavily and lightly impaired. The impairments at D3 were averaged and values bigger than the average are considered heavily impaired, while those with smaller values are lightly impaired. As it is clear in Fig S4, the heavily and lightly impaired olfactory functions have similar slopes of correlation in the PT and PT+PBM group which means that the PBM accelerates the natural recovery after PT with the maintenance of the same trend of olfactory function restoration.

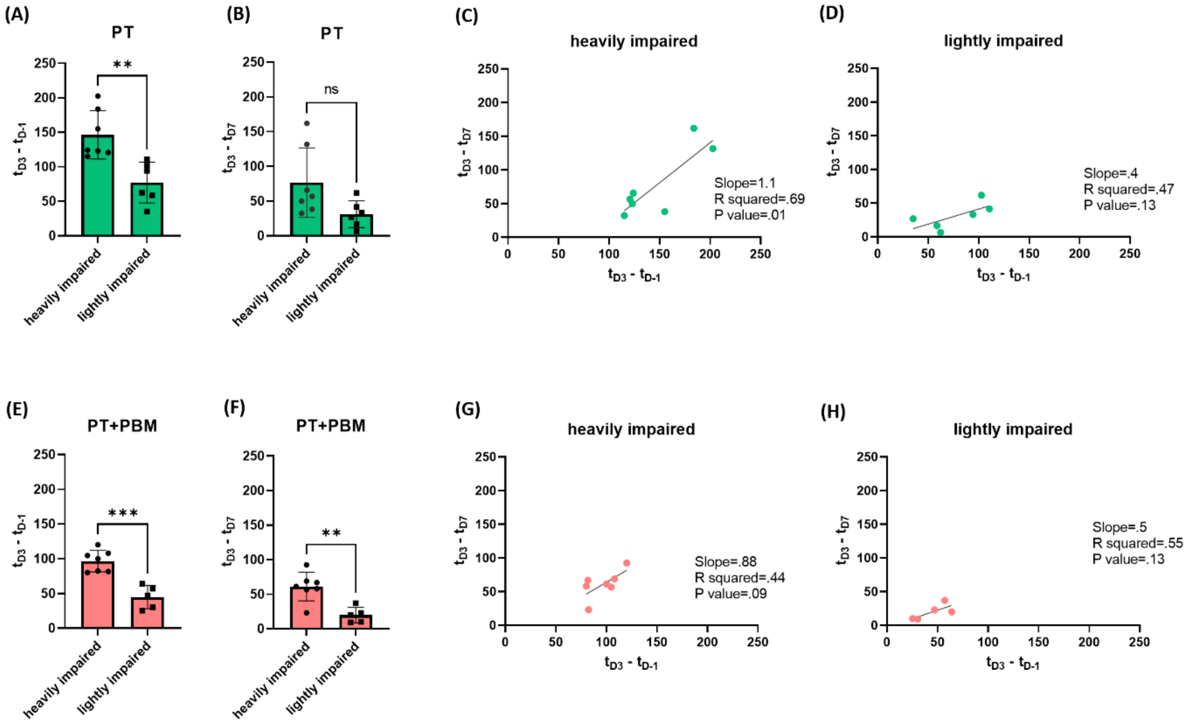


Fig S4. **Correlation between the heavy and light impairment of the olfactory function**. (A), (B) & (E), (F) Comparison between the impairment and restoration of the olfactory function between the heavy and light impairment groups for PT and PT+PBM. (C), (G) correlation between impairment and restoration of the olfactory function for the heavy impairment PT and PT+PBM groups. (D), (H) correlation between impairment and restoration of the olfactory function for the light impairment PT and PT+PBM groups.
